# Programmable microparticles rewire CAR signaling to enable super-physiological expansion of human T cells *in vitro*

**DOI:** 10.1101/2025.07.17.665438

**Authors:** Qinghe Zeng, Landon Flemming, Yuanzhou Chen, Thomas Mazumder, Heinz Hammerlindl, Greg M. Allen, Ricardo Almeida, Jasper Z. Williams, Rogelio A. Hernández-López, Justin Eyquem, Chun J. Ye, Wendell A. Lim, Qizhi Tang, Tejal A. Desai, Xiao Huang

**Affiliations:** School of Biomedical Engineering, Science, and Health Systems, Drexel University, Philadelphia, Pennsylvania, USA; Department of Bioengineering and Therapeutic Sciences, University of California, San Francisco, California, USA; Cell Design Institute and Department of Cellular and Molecular Pharmacology, University of California, San Francisco, California, USA; Department of Medicine and Parker Institute for Cancer Immunotherapy, University of California, San Francisco, California, USA; Department of Microbiology and Immunology, University of California, San Francisco, California, USA; Department of Pharmaceutical Chemistry, University of California, San Francisco, California, USA; Department of Bioengineering, Department of Genetics, Stanford University, California, USA; Diabetes Center, University of California, San Francisco, California, USA; School of Engineering, Institute for Biology, Engineering and Medicine, Brown University, Providence, Rhode Island, USA

## Abstract

T cell proliferative capacity and persistence critically determine the therapeutic success of chimeric antigen receptor (CAR) T cells. However, it remains unknown if and how human CAR-T cells can be externally programmed to reach maximal proliferative capacity. Here, we use programmable PLGA microparticles functionalized with CAR-antigens and CD28-costimulatory antibodies (CAREp) to repeatedly stimulate human CD8^+^ CAR-T cells *in vitro*. CAREp-stimulated CAR-T cells expanded continuously for over 100 days—versus ∼30 days with tumor cell stimulation—and achieved up to 10^18^-fold cumulative expansion, greatly surpassing CD3/28-Dynabeads. Early-phase transcriptomic responses— upregulation of DNA repair, cell cycle, telomere maintenance, and mitochondrial pathways—aligned with long-term outcomes: massive proliferation, telomere stability, robust respiration, and preserved progenitor phenotype by single-cell sequencing. Differentiation and exhaustion signals were broadly suppressed. Transient telomerase activity further supported physiologic expansion. These findings demonstrate that nanoscale-controlled extracellular cues can rewire intracellular signaling to drive durable, super-physiological expansion of functional CAR-T cells.

---

Chimeric antigen receptor (CAR) T cell therapies have transformed treatment of relapsed and refractory B-cell malignancies^1^. Despite their clinical success, major challenges remain—most notably, limited T cell persistence after infusion^1–4^ and inconsistent proliferative capacity during *ex vivo* manufacturing and *in vivo* expansion^5–8^. Understanding the conditions that govern CAR-T cell proliferation and persistence is therefore critical for overcoming these challenges. Particularly, how external signals can be structured to support long-term CAR-T cell expansion and function remains poorly defined.

Physiological T cells possess intrinsic capacity for longevity and tightly regulated proliferation^9–12^, unlike current genetic engineering approaches that enforce intrinsic modifications by bypassing critical cell-cycle checkpoints and risk oncogenic transformation^13–18^. Memory T cells, for example, transiently upregulate telomerase activity upon activation to preserve telomere length and prevent senescence during expansion^19–23^. Massive and sustained proliferation has been demonstrated in natural T cells when stimulated with appropriately balanced antigen and costimulatory inputs^9, 11,12^. These findings raise a key question: can CAR-T cells be externally stimulated— without genetic perturbations—to access a similar proliferative and functional potential? Moreover, while CAR-T cells respond acutely to antigens, costimulatory ligands, and cytokines, chronic stimulation often results in dysfunction and exhaustion^4, 24–26^. It remains unclear how the composition and organization of extracellular cues affect intracellular signaling and long-term fate decisions in engineered T cells. A mechanistic framework linking early signaling dynamics to durable cell states is needed to better understand and potentially modulate CAR-T cell behavior.

Here, we hypothesized that precisely structured extracellular signals, delivered via programmable synthetic materials, could reprogram CAR-T cell signaling and reshape fate decisions. We applied our previously established platform using DNA-scaffolded poly(lactic-co-glycolic acid) (PLGA) microparticles^27, 28^ to create CAR-engaging particles (CAREp)—composed of CAR-target antigens and CD28-costimulatory antibodies (a-CD28 Ab) presented at tunable stoichiometries. These microparticles were used to repeatedly stimulate resting human CD8^+^ CAR-T cells *in vitro*. Surprisingly, CAREp induced sustained CAR-T proliferation for over 100 days—reaching up to 10^18^-fold expansion—far exceeding that observed with repeated tumor cell or commercial CD3/28-Dynabeads stimulations. Expanded cells after multiple rounds of CAREp stimulation were profiled in-depth for phenotype, effector function, metabolic function, telomere dynamics, and clonal evolution. In parallel, bulk RNA sequencing was used to compare early-phase signaling between CAREp and tumor stimulation. We found that CAREp uniquely enriched transcriptional programs associated with cell cycle regulation, DNA repair, chromatin remodeling, telomere maintenance, and mitochondria function—features that aligned with durable expansion, stable telomeres, respiratory fitness, and memory-like cell states (**Fig. 1a**). Together, these results offer mechanistic insight into how nanoscale-structured extracellular signals rewire CAR-T cell behavior—revealing that T cell proliferation and fate can be controlled through the physical format of stimulation, and highlighting principles relevant for both basic T cell biology and future translational design.

**Figure 1.**
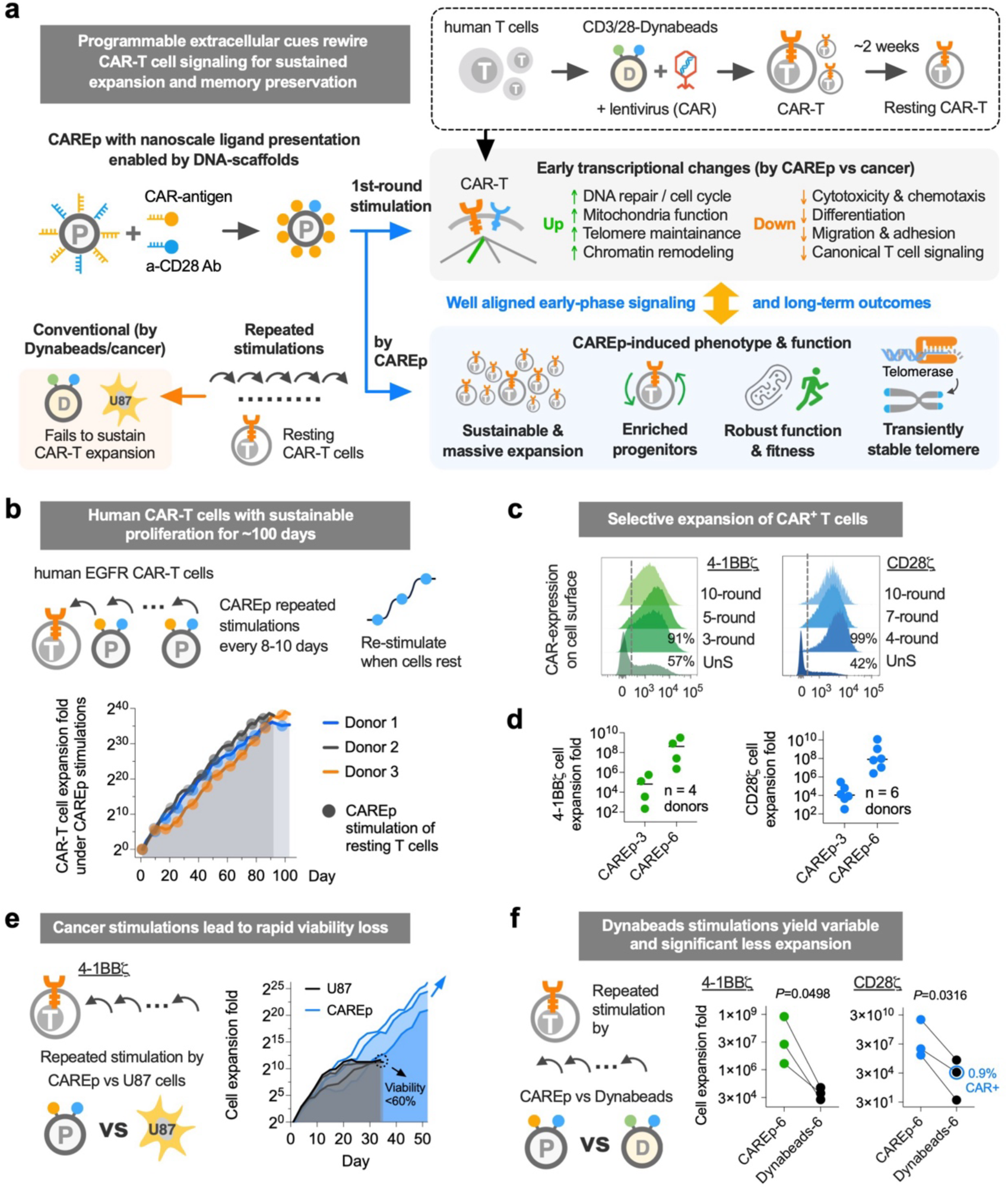
PLGA microparticles functionalized with antigens and CD28 agonist antibodies enabled durable and massive EGFR CAR-T cell expansion *in vitro*. (**a**) Schematic of the CAREp platform and experimental workflow. Human T cells were engineered with conventional CAR lentiviral methods and rested prior to stimulation. DNA-scaffolded PLGA microparticles were functionalized with CAR antigens and CD28 agonists at tunable ratios to create CAR-engaging particles (CAREp). Compared to conventional stimulation by tumor cells, CAREp rewired acute T cell signaling— upregulating pathways for proliferation, metabolism, and telomere maintenance while downregulating canonical differentiation and exhaustion signaling. These rewired early signals aligned with long-term functional outcomes, including massive expansion, memory-like progenitor enrichment, robust metabolic fitness, and transiently stabilized telomere length, demonstrating CAR-T fate and function governed by precise extracellular cues. (**b**) Cumulative expansion of human CD8^+^ EGFR 4-1BBζ CAR-T cells among three donors under repeated CAREp stimulations every 8-10 days when cells are rested from the previous stimulation. The fold of expansion refers to the yielded number of T cells relative to the number present at the time of initial CAREp stimulation. (**c**) Representative 4-1BBζ and CD28ζ CAR surface expression by Myc-tag staining on the expanded CD8^+^ T cells after repeated CAREp stimulations (from n = 3 donors). (**d**) Cumulative expansion folds of CD8^+^ 4-1BBζ-CAR (from n = 4 donors) and CD28ζ-CAR (from n = 6 donors) T cells after 3 and 6 rounds of CAREp stimulation. (**e**) Cumulative expansion of CD8^+^ 4-1BBζ-CAR T cells repeatedly stimulated by a glioblastoma cell line U87 supplemented at 1:1 cancer-to-T cell ratio versus by CAREp when cells are rested from the previous stimulation. (**f**) Cumulative expansion of 4- 1BBζ and CD28ζ CAR-T cells stimulated by 6 rounds of CAREp versus CD3/28- Dynabeads when cells are rested from the previous stimulation. *P* values were determined by two-tailed paired *t*-test. Data in **e**, **f** are from n = 3 healthy donors.

## Massive and sustained expansion of human CAR-T cells from CAREp-stimulations

Agonist antibody against CD28 receptor can provide essential costimulatory signaling for T cell survival and proliferation^29, 30^. To test if the coating of EGFR (epidermal growth factor receptor) antigens on PLGA particles can stimulate CAR-T cell proliferation and if the co-presentation of a-CD28 Ab can augment the activation, we engineered micron-sized CAREp particles with different ratios of the two signals (at 1:0, 9:1, 3:1, 1:3, and 0:1) on the surfaces controlled by the DNA scaffolds (**Fig. 1a** and **supplementary Fig. 1a,b**). Both 2^nd^-generation CAR constructs with a-EGFR nanobody (7D12), CD8α transmembrane domain, and with either 4-1BBζ or CD28ζ costimulatory domain were transduced via lentiviral vectors into human CD4^+^ or CD8^+^ T cells separately isolated from the peripheral blood of healthy donors after CD3/28- Dynabeads (Thermofisher) activation (**Fig. 1a**). The CAR expression was controlled under a constitutive SFFV promoter. Resting T cells from CD3/28-Dynabeads activation and CAR transduction were stimulated with CAREp at 2.5 particles per cell with the supplementation of 30 IU/mL IL-2 in the medium and were maintained at a concentration between 0.5-1.5 million per mL during the *in vitro* culture. Across three donors, CAREp stimulation induced a bell-shaped response pattern for cell proliferation with respect to the EGFR:a-CD28 Ab ligand ratio. The [9:1] ratio consistently yielded the highest cell expansion, while higher or lower proportions of a-CD28-Ab (e.g., [1:3] or [1:0]) led to reduced proliferation (**Supplementary Fig. 1c**). The trend was reproducible across donors and between IL-2 and IL-7/15 supplementations, despite inter-donor variation in absolute cell numbers (**Supplementary Fig. 1c**). Meanwhile, CAREp-induced cell proliferation required cytokine supplementation (**Supplementary Fig. 1d**). Alternative supplementation with IL-7 and IL-15 (5 ng/mL for each) also supported proliferation but was less effective than IL-2 (**Supplementary Fig. 1c**).

Using CAREp-[9:1] with 30 IU/mL IL-2 supplemented, we next evaluated repeated stimulation of CAR-T cells *in vitro*, with resting periods between stimulations. Remarkably, CD8^+^ CAR-T cells exhibited sustained proliferation for up to 100 days across three donors (**Fig. 1b**). In contrast, CAREp-[1:0] without a-CD28 Ab failed to support prolonged expansion (**Supplementary Fig. 1e**), highlighting the necessity of dual-signal input. Notably, stimulation every 4 days did not yield expansion, whereas an ∼8-day interval allowing cells resting from the prior stimulation was required to maintain proliferative capacity (**Supplementary Fig. 1f**). Moreover, resting cells gradually lost viability over ∼7 days without re-stimulation, underscoring their reliance on continuous, antigen-driven signaling. Importantly, only CAR-transduced (EGFR-specific) T cells expanded in response to CAREp, while CAR-negative cells failed to proliferate and likely dwindled (**Fig. 1c**), confirming antigen specificity. Expansion was independent of CAR expression levels, indicating robust signaling irrespective of surface density (**Fig. 1c**). This trend was also observed with the CD28ζ construct, where cumulative expansion reached about 10^4^-fold after three CAREp stimulations and 10^8^-fold after six rounds (**Fig. 1d**). Although donor-to-donor variability is a common feature in human T cell studies^6, 8^, CAREp stimulation appeared to unify proliferative behavior across individuals (**Fig. 1b**). Depending on construct and donor, proliferation sustainability ranged from 90 to 130 days, with cumulative expansion from ∼2^30^ (10^9^) to 2^60^ (10^18^)-fold (**Fig. 1b** and **Supplementary Fig. 1h,i**). These values refer to fold change relative to the starting population at the first CAREp stimulation. When including the preceding 100-1000-fold expansion from standard engineering steps (CD3/28-Dynabeads stimulation and CAR transduction, **Fig. 1a**), the data emphasize the extraordinary proliferative capacity of CAR-T cells when exposed to optimally structured signals.

Interestingly, although CD28ζ-CAR showed a similar pattern of cell expansion after one round of stimulation using particles with different ligand ratios (**Supplementary Fig. 1c,g**), the additional amount of a-CD28 Ab on CAREp-[9:1] compared to CAREp- [1:0], ideal for 4-1BBζ-CAR, was not necessary for CD28ζ-CAR proliferation (**Supplementary Fig. 1g,h**). This is possibly due to the already incorporated CD28 costimulatory signaling domain in the CAR construct. However, CD28ζ-CAR T cells from the same set of donors showed heterogeneous expansion patterns during repeated stimulations with CAREp-[1:0] (**Supplementary Fig. 1i**). For donor 3 that did not respond well to CAREp-[1:0] and a fourth donor (donor 4), a low level of a-CD28 Ab on CAREp-[9:1] enhanced and sustained cell proliferation (**Supplementary Fig. 1j**). Additionally, CD4^+^ T cells showed similar robust proliferation by repeated CAREp stimulations (**Supplementary Fig. 1k,l**).

Next, we asked whether the sustained proliferation observed with CAREp could be replicated using conventional stimulation approaches, including antigen-positive tumor cells and CD3/28-Dynabeads^11, 25, 31^. To account for potential overstimulation effects (**Supplementary Fig. 1f**), T cells were re-stimulated only after a rest phase, mirroring the CAREp protocol. When 4-1BBζ-CAR-T cells were co-cultured with EGFR^+^ glioblastoma U87 cells (1:1 ratio), U87 cells were rapidly eliminated within 2-3 days, followed by T cell expansion for 5-7 days. However, proliferation failed to sustain beyond 3-4 stimulation rounds across all three donors tested (**Fig. 1e**), indicating premature exhaustion or cell death under repeated natural antigen exposure. CD28ζ- CAR T cells from donor 1 maintained proliferation for up to 60 days, but cells from other donors declined after ∼20 days (**Supplementary Fig. 1m**). To further assess antigen format effects, CD28ζ CAR-T cells were stimulated with EGFR- overexpressing K562 leukemia cells. Strikingly, even a single K562 stimulation induced high expression of PD-1, LAG-3, and TIM-3, along with increased Caspase 3/7-activity, in contrast to CAREp-expanded cells (**Supplementary Fig. 1n**). These results suggest that the chemiophysical nature of antigen presentation critically influences T cell fate. We then evaluated CD3/28-Dynabeads in the same rest-stimulate assay. Proliferation varied markedly among donors, and overall expansion after six rounds was orders of magnitude lower than with CAREp stimulation (**Fig. 1f**). Furthermore, Dynabeads do not selectively stimulate CAR^+^ T cells, leading to population dilution. In some donor, CAR^+^ cells were entirely lost (**Fig. 1f**), indicating that standard CD3/28-Dynabeads activation not only underperforms but also expands non-transduced bystanders—underscoring the unique CAR-selective expansion by CAREp.

## Preservation of progenitor-like phenotype during CAREp-mediated expansion

To evaluate how repeated CAREp stimulation affects T cell differentiation, we profiled CAR-T cell phenotype across multiple stimulation rounds using flow cytometry. An initial 10-color panel measured canonical markers of memory and exhaustion, including CD62L, CD45RA, CCR7, CD127, PD-1, LAG-3, and TIM-3. Consistent with proliferation patterns (**Fig. 2a**), CAREp-[9:1] induced the highest percentage of CD62L⁺CD45RA⁺ cells across donors, outperforming suboptimal ratios such as [1:0] or [1:3] (**Fig. 2b,c**). These cells are broadly defined as memory progenitors—subsets associated with enhanced persistence and clinical efficacy^32–37^. For both 4-1BBζ and CD28ζ CAR constructs, the memory progenitor population (CD62L⁺CD127⁺ or CD62L⁺CD45RA⁺) remained stable across up to six stimulations using optimally configured CAREp. Remarkably, even after expansion on the order of 10^6^–10^12^-fold, progenitor-like cells were maintained at ∼20% in 4-1BBζ CAR-T and ∼60% in CD28ζ CAR-T, comparable to the baseline prior to CAREp stimulation (UnS) (**Fig. 2d,e**). This population declined only after eight or more stimulation rounds (CAREp-8 or CAREp-9, **Fig. 2d,e**), suggesting a late-stage shift toward terminal differentiation.

**Figure 2.**
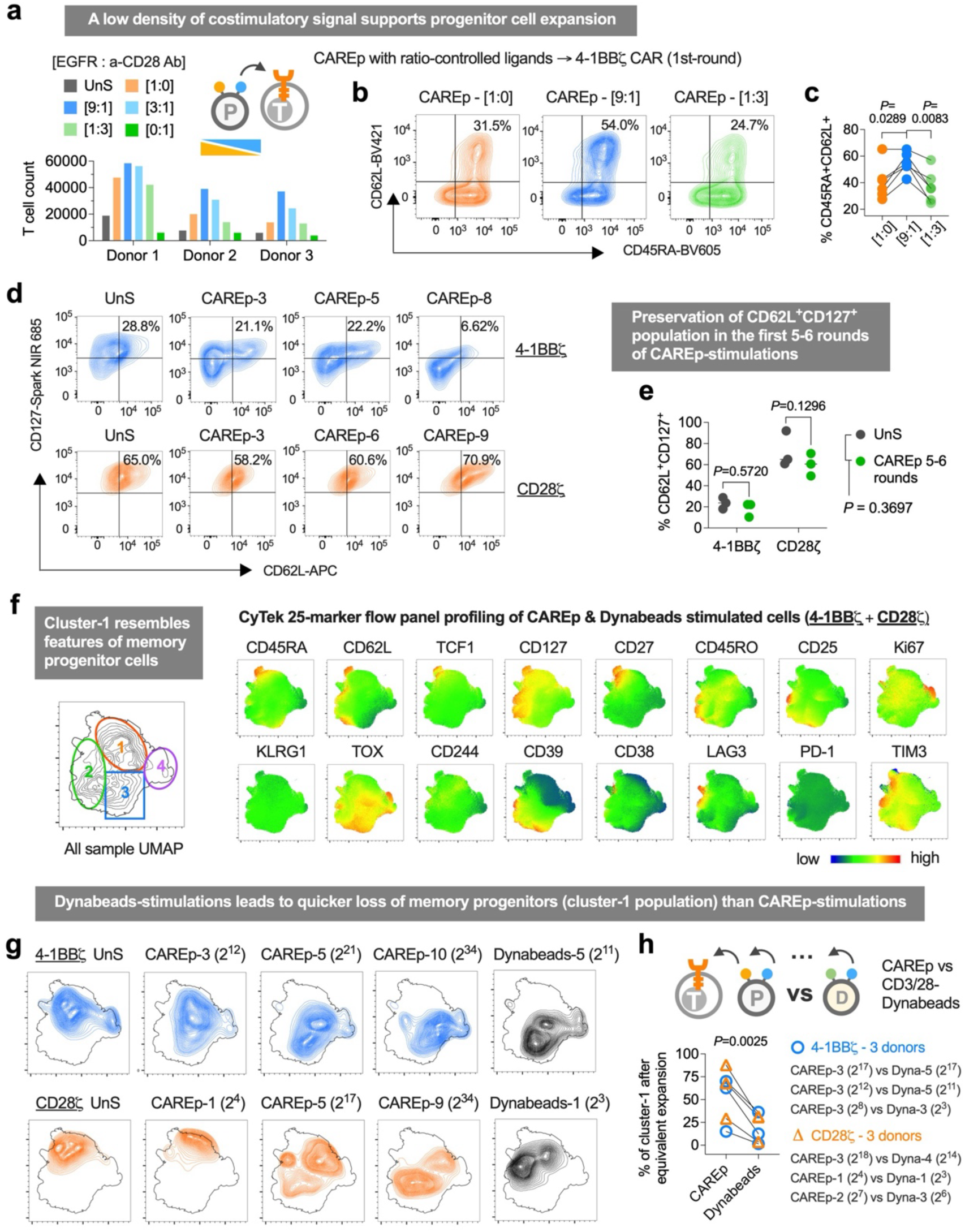
Memory-like progenitors are better preserved during repeated stimulations by CAREp versus CD3/28-Dynabeads. (**a**) Relative expansion of CD8^+^ 4-1BBζ CAR-T cells 10 days after the first stimulation by CAREp coated with varying densities of EGFR-antigens and a-CD28 agonist antibodies (a-CD28 Ab) ([1:0], [9:1], [3:1], [1:3], [0:1]) in the presence of 30 IU/mL IL-2. UnS: unstimulated cells. (**b**) Representative CD62L and CD45RA expression of the expanded CD8^+^ T cells at 10 days after the first CAREp ([1:0], [9:1], [1:3]) stimulation. (**c**) CD45RA^+^CD62L^+^ population in CAREp ([1:0], [9:1], [1:3])-expanded cells in the presence of 30 IU/mL IL-2 or IL-7/15 (5 ng/mL each). Data are from n = 6 independent samples, and *P* values were determined by one-way ANOVA with Tukey’s multiple comparisons test. (**d**) Representative flow plots of CD127 and CD62L expression on CD8^+^ 4-1BBζ and CD28ζ CAR T cells that experienced 0-9 rounds of CAREp stimulation. (**e**) CD62L^+^CD127^+^ population of CD8^+^ EGFR-CAR T cells (4-1BBζ and CD28ζ constructs) before (UnS) and after 5-6 rounds of CAREp stimulation. Data represent mean of n = 3 donors, and *P* values were determined by two-way ANOVA with Sidak’s multiple comparisons. (**f**) UMAP and key marker expression of the pooled CD8^+^ EGFR-CAR T cell samples from repeated CAREp or CD3/28-Dynabeads stimulations after the immunostaining using a 25-marker spectrum flow panel. (**g**) Distribution of the expanded cells (with expansion fold in the bracelet) by Dynabeads or CAREp stimulations on the pooled UMAP in **f**. (**h**) Cluster-1 population in CAREp or CD3/28-Dynabeads stimulated 4-1BBζ or xCD28ζ CAR T cells with equivalent folds of cell expansion. Data are from n = 3 donors, and *P* value was determined by two-tailed paired *t*-test. Samples in **a-h** are from n = 3 healthy donors.

To gain a deeper insight into cell phenotypic changes, we applied spectrum flow cytometry using a 25-marker panel including CD45RA, CD62L, TCF1, CD127, CD27, CD28, CCR7, CD45RO, CD95, CD25, Ki67, ID2, CXCR3, HELIOS, IRF4, PRDM1, HLA-DR, KLRG1, TOX, CD244, CD39, CD38, LAG3, PD-1, TIM3. Cells from multiple CAREp and CD3/28-Dynabeads stimulations were pooled and visualized using a UMAP projection (**Fig. 2f** and **Supplementary Fig. 2a**). Due to inconsistent expansion with Dynabeads, only cells from ≤5 rounds could be included. Manual gating based on memory- and exhaustion-associated markers resolved four clusters. Cluster-1, characterized by high CD62L, CD27, CD45RA, and CD127 and low PD-1, TIM-3, LAG-3, and CD39, was defined as a memory progenitor-like state (**Fig. 2f** and **Supplementary Fig. 2b**). We observed a gradual phenotypic progression from cluster-1 to more differentiated states with repeated CAREp stimulation, with divergence in trajectories between the two CAR constructs (**Fig. 2g** and **Supplementary Fig. 2c-f**). Importantly, cluster-1 remained preserved through 3-5 CAREp rounds, consistent with 10-color panel data (**Fig. 2d,e**). When comparing CAREp and Dynabeads stimulations matched for expansion fold, cluster-1 frequencies were consistently more than two folds higher with CAREp in both CAR constructs (**Fig. 2g,h** and **Supplementary Fig. 2c,e,g, and h**), suggesting a quicker differentiation induced by Dynabeads. Collectively, these results demonstrate that CAREp stimulation enables massive expansion while delaying terminal differentiation, maintaining a progenitor-like phenotype through the early-to-mid phases of *in vitro* proliferation (up to rounds 5-6), during which ∼10^4^–10^8^-fold expansion was achieved.

## Decoupled T cell signaling and telomerase regulation

When T cells encounter antigen-expressing cancer targets, canonical signaling pathways—including calcineurin-NFAT, PKCθ-NF-κB, Ras-Raf-Erk, and chemotaxis pathways—are activated to mediate functions such as migration, adhesion, and cytotoxicity^38^. However, in CAR-engineered T cells, chronic antigen exposure often leads to rapid exhaustion and functional decline^4, 25^. To investigate how signaling profiles differ between stimulatory formats that do or do not support long-term CAR-T proliferation, we stimulated CD28ζ CAR-T cells from three donors with either CAREp-[1:0] or EGFR^+^ U87 tumor cells and performed bulk RNA sequencing at 24 hours. Unstimulated CAR-T cells (UnS) served as controls. For U87 co-cultures, CD8+ T cells were FACS-purified to exclude tumor cell transcripts. Principal component (PC) analysis revealed that CAREp and U87 induced distinct transcriptomic programs, overriding inter-donor variation (**Fig. 3a** and **Supplementary Fig. 3a**). CAREp stimulation upregulated pro-survival and metabolic genes such as CD27, BCL2L10, MT-RNR1 and CYP1B1, while downregulating effector and exhaustion-associated transcripts, including PDCD1, LAG3, TNFRSF18, IL23A, and IL12RB2 (**Fig. 3b**).

**Figure 3.**
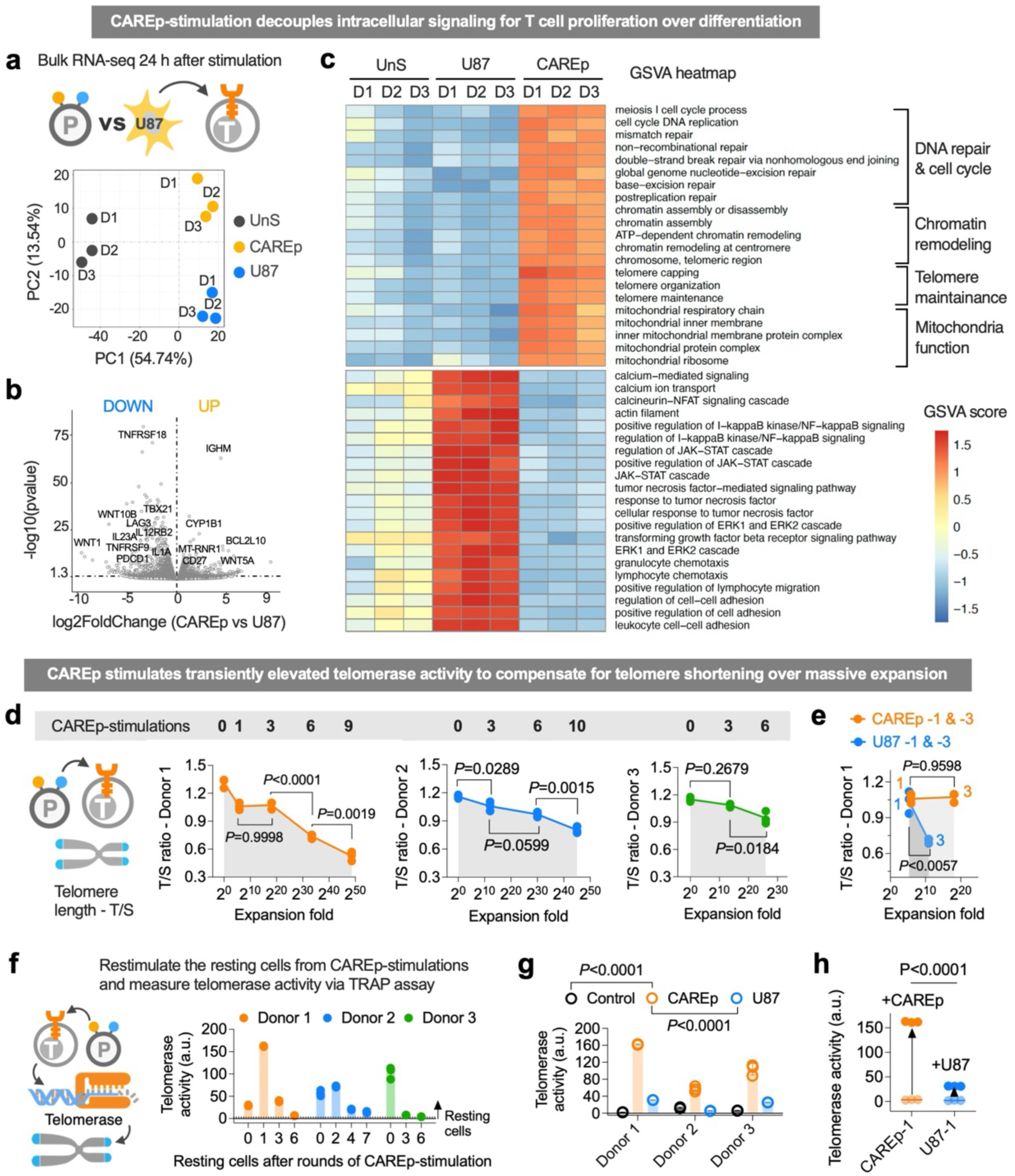
CAREp rewired gene program supporting proliferation over differentiation and induced upregulated telomerase activity to stabilize telomere length. (**a**) Principal component analysis (PCA) of CD8^+^ CD28ζ CAR-T cell transcriptome by bulk RNA sequencing at 24 hours after CAREp or U87 stimulation or CAREp-unstimulated cells. Data are from n = 3 donors (D1-3). (**b**) Volcano plots of differentially expressed genes between CAREp and U87 stimulated cells in **a**. (**c**) GSVA of induced signaling and pathways that are significantly different between CAREp and U87 stimulated cells (adjusted *P* value < 0.01). (**d**) Telomere lengths of the expanded CD8^+^ CD28ζ CAR-T cells after varying rounds of CAREp stimulation by the T/S assay. Data are mean ± SD from n = 3 technical replicates, and *P* values were determined by one-way ANOVA with Tukey’s multiple comparisons test. (**e**) Telomere length of CD8^+^ CD28ζ CAR-T cells after 1 and 3 rounds of CAREp versus U87 stimulations. Data are mean ± SD from n = 3 technical replicates, and *P* values were determined by two-way ANOVA with Sidak’s multiple comparisons test. (**f**) Telomerase activity by the TRAP analysis of CD8^+^ CD28ζ CAR-T cells at 3 days after CAREp-restimulation of resting cells from varying rounds of CAREp stimulation. Data are mean ± SD from n = 3 technical replicates. (**g**) Telomerase activity of CD8^+^ CD28ζ CAR-T cells at 3 days after CAREp versus U87 stimulation. Donor 1 was measured at the second round of CAREp or U87 stimulation, and donors 2 and 3 were measured at the first round of CAREp or U87 stimulation. Data are mean ± SD from n = 3 technical replicates, and *P* values were determined by two-way ANOVA with Tukey’s multiple comparison test. (**h**) Telomerase activity at 3 days after the restimulation of CD8^+^ CD28ζ CAR-T cells that experienced 1-round of CAREp or U87 stimulation, compared to the unstimulated controls. Data are mean from n = 3 technical replicates, and *P* value was determined by two-tailed unpaired *t*-test.

Through gene ontology (GO) enrichment analysis, pathways (zone 2) including the apoptotic process, TGF beta signaling, actin cytoskeleton, and inflammatory response were significantly upregulated by the U87 stimulation but remained low by CAREp stimulation, compared to the unstimulated group (**Supplementary Fig. 3b**). In contrast, important processes associated with cell proliferation (zone 3 and zone 4), such as DNA repair, cell cycle, cytoplasmic translation, protein transport, chromatin organization & remodeling, telomere maintenance, mitochondria translation, response to oxidative stress, were significantly upregulated by CAREp stimulation, but were less upregulated (zone 4) or stayed low (zone 3) by U87 stimulation (**Supplementary Fig. 3b**). Interestingly, cell signaling associated with signal transduction, cell migration, and chemotaxis remained high for U87 stimulation compared to the unstimulated group, but they were downregulated by CAREp stimulation (**Supplementary Fig. 3b**).

To further identify the signaling and processes of GO terms that are apparently up- or down- regulated between CAREp and U87 treatments, we next run gene set enrichment analysis (GSEA) and narrowed down the list, although quite a few of them showed a low P-value but high FDR q-val (**Supplementary Fig. 3c**). To obtain a quantitative view of these pathways among all three treatments, gene set variation analysis (GSVA) was run among all samples (**Fig. 3c**). Strikingly, the pathways associated with DNA repair, chromatin remodeling, telomere maintenance, and mitochondria function were consistently upregulated, while those signaling for Calcineurin-NFAT, PKC-θ-NF-κB, Ras-Raf-Erk, JAK-STAT cascades, and chemotaxis were kept low or downregulated by CAREp among donors, compared to T cells stimulated by U87 cells (**Fig. 3c**). This demonstrated that pro-survival and proliferation pathways (DNA repair, cell cycle, chromatin remodeling, telomere maintenance, oxidative stress response) were markedly enriched in CAREp-stimulated cells, whereas effector function pathways (inflammatory response, NFκB, JAK-STAT, calcium/NFAT signaling, chemotaxis/adhesion) were relatively subdued compared to tumor-stimulated cells, indicating the rewired gene program favoring proliferation over differentiation, which corresponds well with the sustainable proliferation through repeated stimulations.

To investigate whether these signaling differences translate to differential telomere dynamics, we measured telomere length across multiple rounds of CAREp stimulation using quantitative PCR (T/S assay)^39^ (**Fig. 3d**). Like other physiological cells, continuous expansion yielded a gradually decreased telomere length among all three donors (**Fig. 3d** and **Supplementary Fig. 3d,e**). However, in all donors, telomere length remained stable within the first three stimulations (approximately 20 cell divisions) (**Fig. 3d,e** and **Supplementary Fig. 3f**). Among the three donors, donor 1 showed the best proliferative sustainability under U87 repeated stimulations (**Fig. Supplementary 1m**). Therefore, we were able to freeze-stock enough cells and analyze the telomere length after three rounds of stimulations. Besides less overall expansion fold (∼10 divisions by U87 vs 20 divisions by CAREp), cells yielded from three U87 stimulations showed significantly decreased telomere length compared to one round of stimulation, whereas telomere length stayed the same between one and three rounds of CAREp stimulation (**Fig. 3e**). This further indicates that CAREp stimulation preserves telomere ends during early massive expansion, whereas accelerated telomere erosion happens with cancer-stimulated cell expansion.

To maintain self-renewal capacity and super longevity, memory T cells can upregulate their telomerase activity upon activation to compensate for telomere shortening during cell proliferation^19–21^. To investigate the telomerase activity after CAREp stimulations, frozen-stocked resting cells from multiple rounds of stimulation were thawed, stimulated with CAREp, and analyzed after three days using telomerase repeated amplification protocol (TRAP)^40^. As expected, resting cells have a neglectable level of telomerase activity (**Supplementary Fig. 3g**), but early CAREp-stimulated cells (rounds 0-2) displayed strong telomerase induction (**Fig. 3f**). Notably, this inducibility declined in later rounds, consistent with progressive loss of stemness. As a comparison, U87-stimulated CAR-T cells showed significantly lower telomerase activity across all donors (**Fig. 3g**). Consistent with the above-mentioned telomere length retention by initial CAREp stimulations but telomere erosion by initial U87 stimulations (**Fig. 3e**), a second round of CAREp stimulation induced a significantly higher level of telomerase activity than a second round of U87 stimulation (**Fig. 3h**).

Together, these results show that CAREp stimulation induces a distinct transcriptional state that temporally enhances telomerase activity and supports telomere integrity during the early stages of massive proliferation. This proliferative capacity aligns with elevated signaling in DNA repair, chromatin remodeling, and mitochondrial regulation, and occurs in the absence of typical effector or exhaustion-inducing signals— demonstrating that CAREp rewires CAR-T cells toward a physiologically regulated expansion program.

## Mitochondria fitness and effector function of the expanded cells

Given the upregulation of mitochondria-associated pathways in CAREp-stimulated CAR-T cells (**Fig. 3c**), we next assessed mitochondria fitness using the Seahorse MitoStress assay^41, 42^. CD8^+^ CAR-T cells rested after repeated CAREp stimulation were sequentially treated with respiration inhibitors—oligomycin, FCCP, and rotenone/antimycin A—to measure oxygen consumption rate (OCR) as a readout of respiratory function (**Fig. 4a**). For both 4-1BBζ-CAR and CD28ζ-CAR constructs, cells expanded for 3-6 CAREp rounds retained or improved respiratory capacity compared to baseline (**Fig. 4a** and **Supplementary Fig. 4a**). Across multiple donors, we observed a consistent increase in spare respiratory capacity (SRC), supporting enhanced mitochondrial performance (**Fig. 4b**). These functional findings align with transcriptomic data indicating CAREp stimulation promotes mitochondrial function and fitness (**Fig. 3c**).

**Figure 4.**
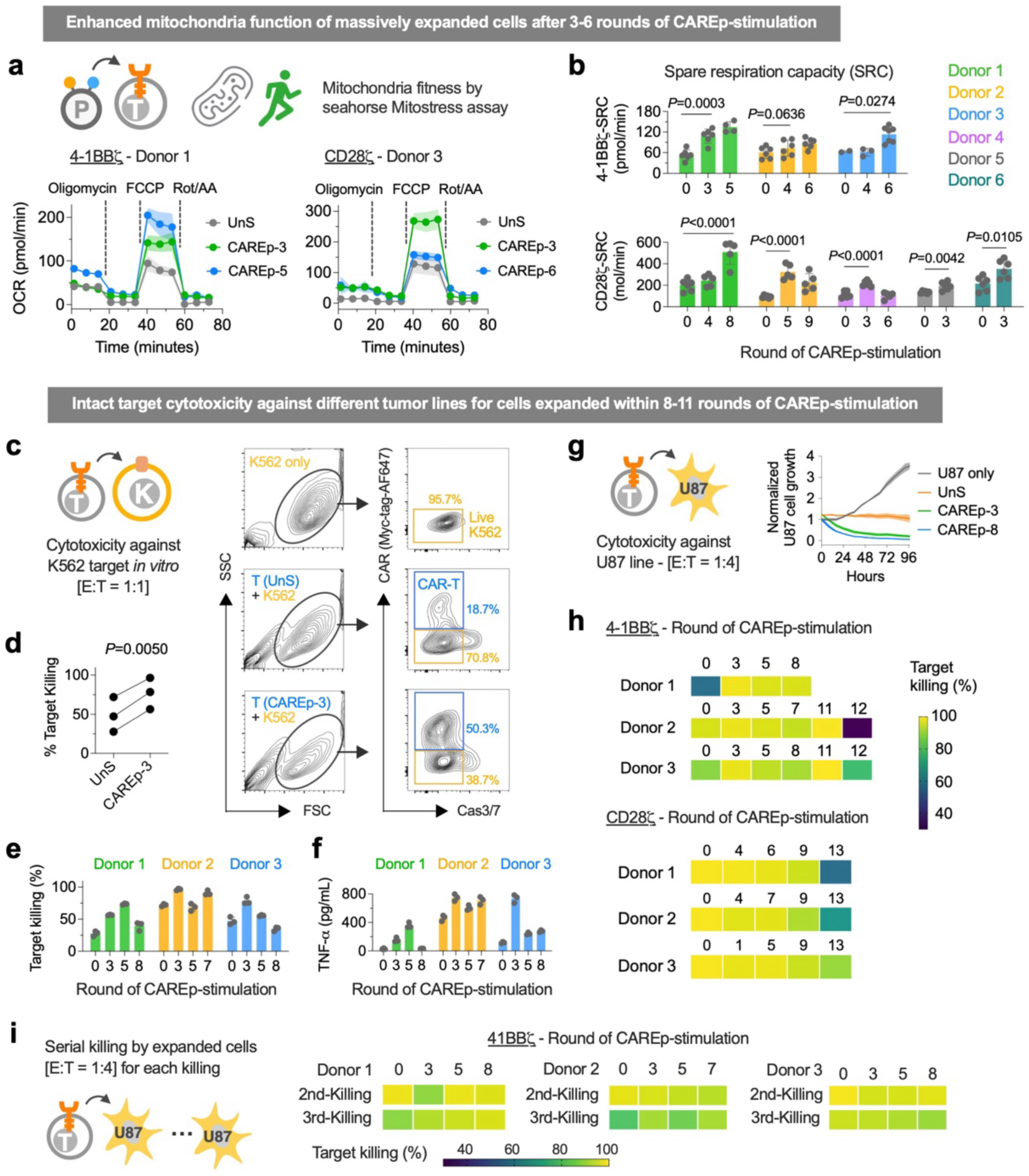
Massively expanded cells demonstrate robust mitochondria fitness and effector function. (**a**) Representative oxygen consumption rate (OCR) of CAREp- expanded CD8^+^ 4-1BBζ and CD28ζ CAR-T cells treated with respiration inhibitors (Oligomycin, FCCP, Rot/AA) from Seahorse Mitostress assay. Data are mean ± SD from n = 6 biological replicates. (**b**) Spare respiration capacity (SRC) of CAREp- expanded CD8^+^ 4-1BBζ and CD28ζ CAR-T cells from Seahorse Mitostress assay. Data are mean ± SD from n = 6 biological replicates. (**c**) Flow cytometry plot of EGFR- overexpressing K562 cells at 4 days after the co-culture with CD8^+^ 4-1BBζ CAR-T cells with or without CAREp-stimulation. Effector-to-target (E:T) ratio was controlled at 1:1. Non-apoptotic K562 cells (yellow gates), distinguished from live CAR^+^ T cells (blue gates), were analyzed for target killing efficacy compared to K562 only control. (**d**) Target killing efficacy (%) between unstimulated CAR-T cells and 3-rounds of CAREp-stimulated cells in **c.** Data are from n = 3 donors, and *P* value was determined by two-tailed paired *t*-test. (**e-f**) Target killing efficacy (%) at day-4 (**e**) and supernatant TNF-α level at day-2 (**f**) after the co-culture of K562 and T cells experienced 0-8 rounds of CAREp-stimulation. Data are mean ± SD from n = 3 biological replicates. (**g**) Growth of U87 cell line co-cultured with CD8^+^ 4-1BBζ CAR-T cells expanded after 0 (UnS), 3, and 8 rounds of CAREp stimulation. Data are mean ± SD from n = 4 biological replicates. (**h**) Target killing efficacy (%) of CAREp-expanded CD8^+^ 4-1BBζ and CD28ζ CAR-T cells against U87 cell line at 72 hours after the co-culture. Data are mean from n = 4 biological replicates. E:T ratio is controlled at 1:4. (**i**) Serial killing of CAREp-expanded CD8^+^ 4-1BBζ CAR-T cells against U87 cells for three rounds. E:T ratio is controlled at 1:4 for each round. Data are mean from n = 4 biological replicates.

To assess effector function, we performed tumor cell killing assays using K562 cells engineered to express truncated EGFR. This target was relatively resistant to CAR-T cytotoxicity, particularly for CD28ζ CAR-T cells, which underwent apoptosis upon co-culture and failed to eliminate the target (**Supplementary Fig. 1n**). In contrast, 4-1BBζ CAR-T cells exhibited modest killing ability, which was markedly enhanced after three CAREp stimulations (**Fig. 4c,d**). These expanded cells also persisted at higher numbers following a 4-day co-culture (**Fig. 4c**). However, this cytotoxic advantage diminished after 7-8 stimulation rounds (**Fig. 4e**), indicating a functional ceiling after extended expansion. TNF-α secretion patterns mirrored killing efficacy, peaking after three CAREp rounds and declining after extensive expansion (**Fig. 4f**).

To evaluate killing of a more physiologically relevant target, we co-cultured CAREp-expanded CAR-T cells with EGFR^+^ U87 glioblastoma cells at an effector-to-target (E:T) ratio of 1:4 (**Fig. 4g**). Through an Incucyte imaging assay, CAREp-expanded cells (CAREp-3 and CAREp-8) exhibited more rapid killing kinetics compared to unstimulated controls (UnS) (**Fig. 4g**). Across all donors and both CAR constructs, T cells expanded for up to 11 CAREp rounds maintained potent cytotoxicity (**Fig. 4h**). Beyond 11 rounds, efficacy declined, consistent with the expansion plateau. For 4-1BBζ CAR-T cells, 3 rounds of CAREp expansion yielded cells with improved cytotoxicity, similar as the trend observed in K562 killing (**Fig. 4d,h**). Cytokine secretion during U87 killing was also profiled (**Supplementary Fig. 4b-e**). CAREp- expanded T cells produced similar levels of TNF-α and IL-2 as unstimulated controls at 48 hours post co-culture (E:T = 1:1), but secreted lower levels of IFN-γ (**Supplementary Fig. 4c,e)**. Finally, we evaluated serial killing capacity in repeated co-culture cycles at E:T = 1:4. CAR-T cells expanded through up to 8 rounds of CAREp maintained robust serial killing over three to four tumor challenges (**Fig. 4i** and **Supplementary Fig. 4f**). Given that extensively stimulated cells showed potent cytotoxicity and serial killing efficacy, lower level of IFN-γ secretion may be due to the cell population evolution with contracted short-lived effector cells and enriched memory cells (**Fig. 2**). Notably, IFN-γ signaling is dispensable—and potentially inhibitory—for CD8⁺ T cell memory formation^43–45^.

## Clonal enrichment of memory progenitor and durable effector states

To dissect the phenotypic and clonal trajectories underlying CAREp-induced expansion, we performed single-cell RNA sequencing (scRNA-seq) with paired CITE-seq and TCR tracking on CD28ζ CAR-T cells across four timepoints per donor (n = 3 donors; 12 total samples; **Fig. 5a**). Resting cells were barcoded with hashtags, pooled, and stained with a 130-marker CITE panel prior to droplet-based sequencing. This strategy allowed for integrated transcriptomic, surface protein, and clonal analysis while minimizing batch effects (**Fig. 5a** and **Supplementary Fig. 5a**). After data quality control, robust numbers of cells were identified among the three libraries, although cells from donor 3 after 5 and 9 rounds of CAREp stimulation showed less robust quality after the freeze-thaw cycle during sample preparation (**Supplementary Fig. 5a**).

**Figure 5.**
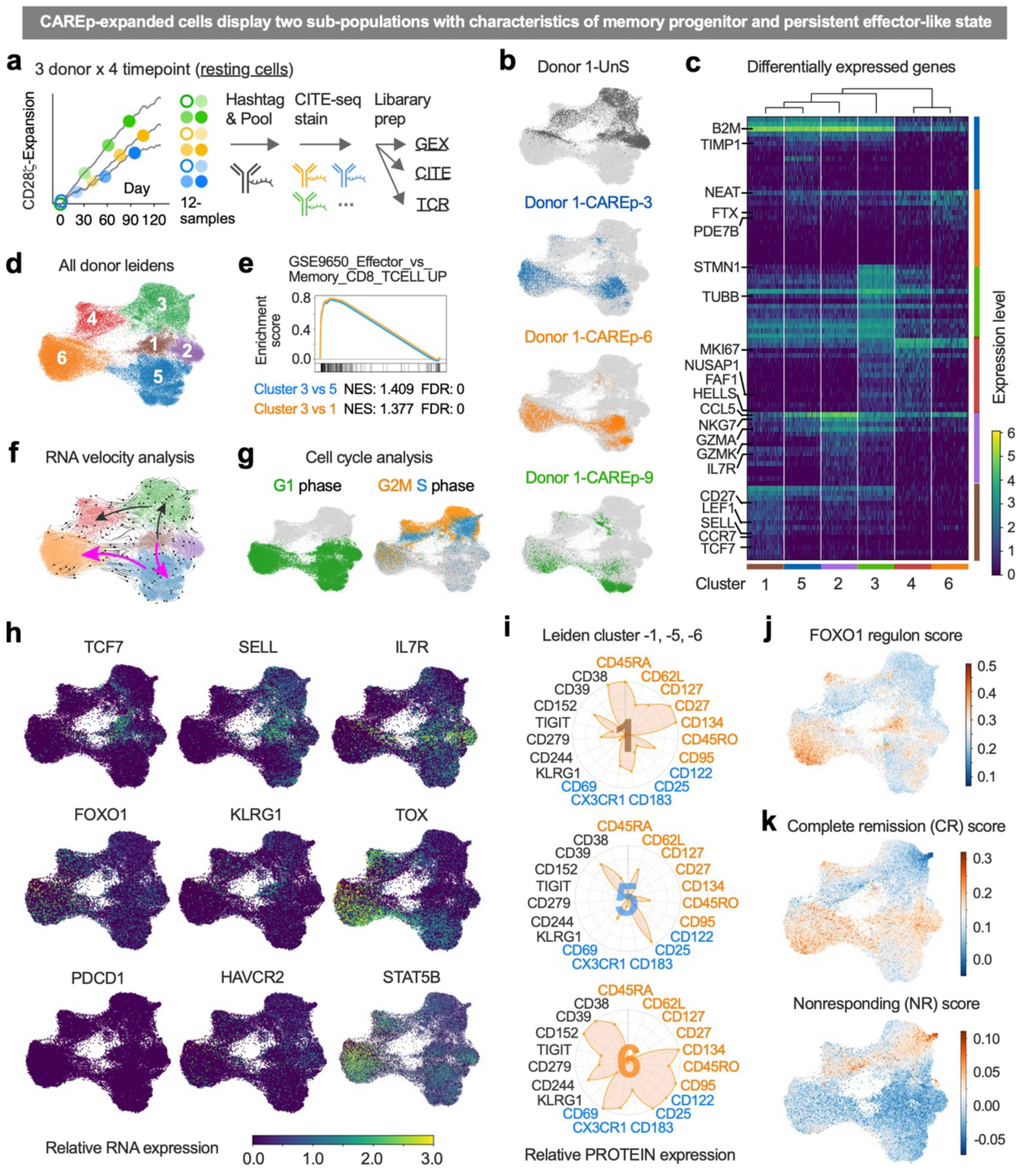
CAREp stimulation generates two transcriptionally distinct T cell subpopulations with memory progenitor and persistent effector-like states. (**a**) Schematic of single cell RNA sequencing of resting CD8^+^ CD28ζ CAR-T cells from 3 donors x 4 timepoints with simultaneous analysis of transcriptome (GEX), 130 surface markers (CITE), and TCR repertoire (TCR). 12 samples (5,000 cells each for UnS, 3000 each for other timepoints) were individually stained with hashtag and pooled together for sequencing. (**b**) UMAP of GEX data highlighting cells from 0, 3, 6, 9 rounds of CAREp stimulation (donor 1). Light grey dots represent cells from all samples. (**c-d**) Unsupervised leiden clustering of GEX data and differentially expressed genes in the leidens. (**e**) Enrichment score of leiden cluster 3 vs 5 and 3 vs 1 from GSE9650 gene set enrichment analysis (GSEA). NES: normalized enrichment score; FDR: false discovery rate. (**f**) Trajectory of cells by RNA velocity analysis of GEX data. (**g**) Cell cycle distribution (G1, G2M, and S) in all sample UMAP from cell cycle analysis. (**h**) Relative RNA expression of selected genes on the GEX-UMAP. (**i**) Relative expression of selected surface markers (protein) in leiden 1, 5, and 6. Markers in orange, blue, and black are respectively associated with memory, effector, and exhaustion phenotypes. (**j-k**) Gene set scoring (Scanpy) of all sample GEX data using reported gene sets that showed correlation with patient complete remission (CR) or nonresponding (NR)^14, 50^.

UMAP projection and Leiden clustering of gene expression data revealed six distinct clusters with gene signatures for memory and effector phenotypes^46, 47^ (**Fig. 5b-d** and **Supplementary Fig. 5b**). Clusters 1-4 contained mostly unstimulated cells; clusters 5 and 6 were enriched with CAREp-expanded cells (**Fig. 5b** and **Supplementary Fig. 5b-g**). Cluster 1 expressed canonical memory progenitor markers (e.g., IL7R, CD27, LEF1, SELL, CCR7, and TCF7), while cluster 3 and cluster 4 respectively showed effector-associated genes (e.g., CCL5, GZMA) and terminal differentiated genes (e.g., TOX, BATF), reflecting conventional T cell differentiation (**Fig. 5c-e** and **Supplementary Fig. 5b-g**). A high similarity was observed between cluster 1 and 5, while cluster 6 showed dramatic difference from these two clusters (**Fig. 5c**). Cluster 5 showed decreased expression of memory-associated genes (TCF7, CD27, LEF1) and increased expression of effector-associated genes (CCL5, GZMA) compared to cluster 1, suggesting a differentiation shift to the effector-like state. Yet, gene set enrichment analysis (GSEA) revealed that cluster 5 still represented a higher level of progenitor state than cluster 3 (**Fig. 5e**). Consistently, RNA velocity analysis showed two distinct trajectories of cell differentiation: one from cluster 1 to 3 to 4 (grey), and the other one from cluster 1 to 5 to 6 (magenta), aligning well with the GSEA results (**Fig. 5e,f**). Thus, CAREp stimulation bifurcated the differentiation trajectory: instead of all cells progressing from Memory (cluster 1) → Effector (cluster 3) → Terminal (cluster 4) as in conventional differentiation, many CAREp-expanded cells entered an alternative route: Memory (cluster 1) → Effector-memory hybrid (cluster 5) → Durable effector-like (cluster 6) (**Fig. 5f**). Interestingly, cell cycle analysis showed that cells at clusters 1, 2, 5, and 6 primarily reside at G1 phase, while cluster 3 and 4 reside at G2M and S phases (**Fig. 5g**), responding to the acutely regulated cell cycle pathways by CAREp stimulation as well (**Fig. 3c**).

Compared with cluster 1 for both RNA and protein expression, cluster 5 maintained moderate levels of memory markers SELL/CD62L (both RNA and protein) and IL7R/CD127 (RNA), upregulated effector marker IL2RA/CD25 (both RNA and protein), and maintained low levels of senescence or exhaustion markers (PDCD1/CD279/PD-1, LAG3, HAVCR2/TIM3, KLRG1, TIGIT, both RNA and protein), indicating an effector-memory state (**Fig. 5h,I** and **Supplementary Fig. 5d-i**). Compared with cluster 5, cluster 6 showed a decrease of SELL/CD62L (both RNA and protein) and IL7R/CD127 (RNA), an increase of effector markers including CD69, CD244, CX3CR1, CD122, CD45RO (protein), and low levels of PDCD1, LAG3, HAVCR2/TIM3, TIGIT, and CD39 (RNA) (**Fig. 5h,I** and **Supplementary Fig. 5d-h**). Although cluster 6 is transcriptomically similar to a terminal effector (like cluster 4) (**Fig. 5c**), it lacked surface expression of KLRG1 and TIGIT, and instead showed persistent CD25, CD122, and STAT5 activation (**Fig. 5h,I** and **Supplementary 5i**). Therefore, cells in cluster 6 clearly resemble the durable effector-like state during chronic antigen exposure^48, 49^. From CAR-T cell clinical trials, several gene sets in manufactured cell products have been identified to importantly correlate with patient remission or non-responding^14, 50^. To further assess the cell quality yielded from CAREp-stimulations, we carried out gene set scoring (Scanpy) of three gene sets including FOXO1 regulon^14^, complete remission (CR), and nonresponding (NR) in the GEX data set^50^. Interestingly, cluster 5 and 6 showed high scores of FOXO1 regulon and CR gene sets and low scores of NR gene set, comparative to memory progenitor cells in cluster 1 (**Fig. 5j,k**).

From TCR repertoire analysis, we identified a clear trend of clonal enrichment over time among all donors, although that CAREp stimulation is TCR-independent (**Fig. 6a**). Due to suboptimal gene counts in clusters 4 and 6, TCR repertoire was more successfully tracked in clusters 1-3 and 5 (**Supplementary Fig. 5c**). Still, clones with drastically different lifespan and proliferative timeline were identified (**Fig. 6b**). For example, some clones (e.g., clones 4 and 28) vanished by day ∼30; some clones (e.g., clone 43, 89, and 39) actively proliferate for 60-70 days and stabilized after 90 days; and some other clones (e.g., clone 14, 47 and 19) actively proliferate beyond 90-100 days (**Fig. 6b**). We then sorted clones with similar patterns of lifespan to three groups (G1 – untraceable after 30 days, G2 – untraceable after 70 days, and G3 – traceable at 90-100 days) to track their phenotypic evolution (**Fig. 6c**). Notably, clones destined to fade early (G1) showed an overlap of memory (CD127, CD62L, and CD27) alongside effector/exhaustion (CX3CR1, KLRG1, CD244, TIGIT, CD279, CD152, CD223) features at the gene/protein level (**Fig. 6d-f**). In contrast, the longer-lived clones (G2 and G3) largely lacked these exhaustion markers from the start, suggesting a more pristine progenitor state (**Fig. 6d-f**). Similarly, between G2 and G3 and among clones of different lifespan, the expression of effector and exhaustion associated markers (CCL5, TOX, CD38, CD25, CD45RO) as well as the levels of T cell activation processes at a similar timeframe were relatively lower in longer-living clones (**Fig. 6d-f** and **Supplementary Fig. 6a-d**).

**Figure 6.**
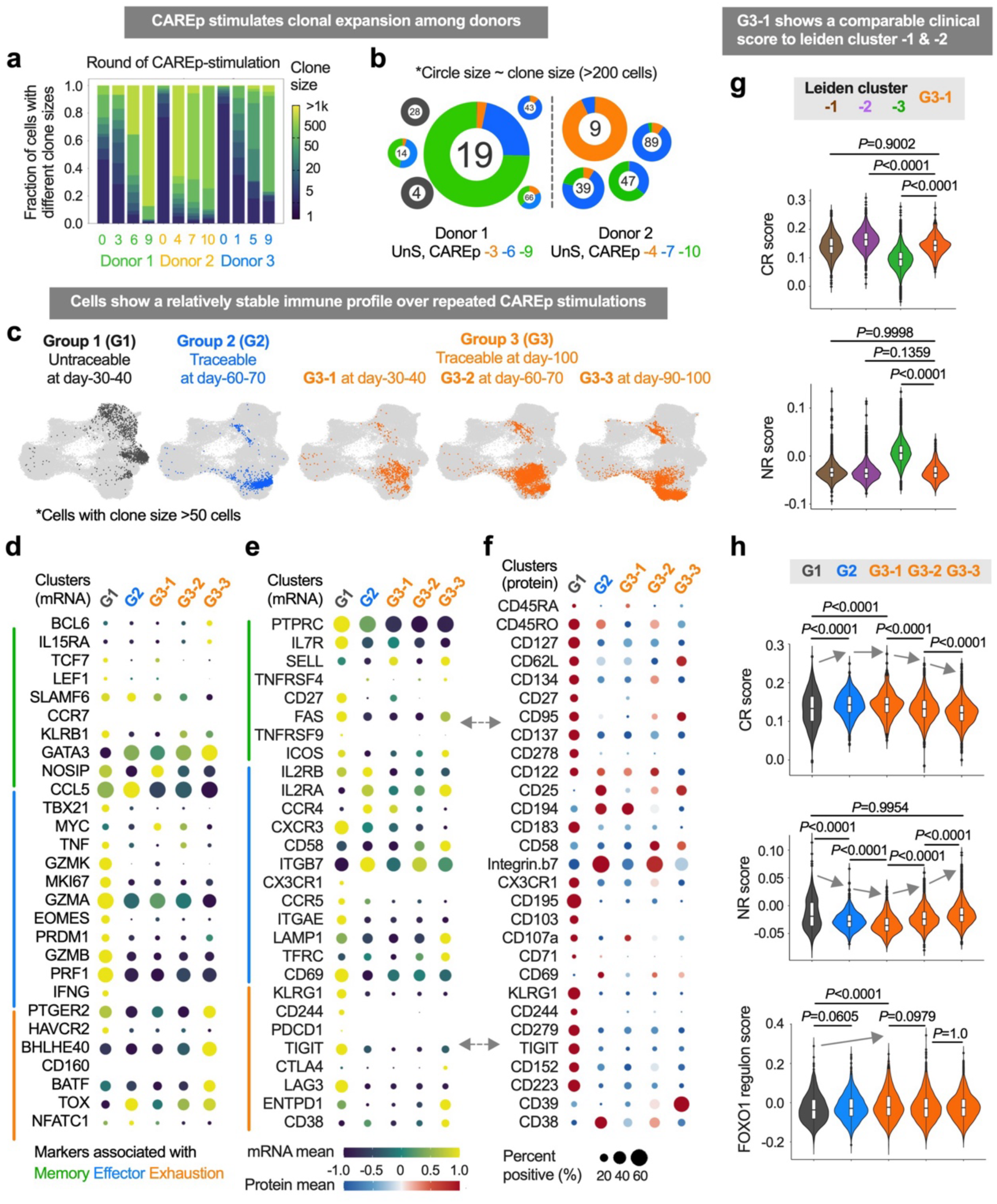
Clonal tracking reveals heterogeneous proliferative lifespans, with CAREp- expanded long-lived clones maintaining stable, progenitor-like states. (**a**) Clonal distribution of CD8^+^ CD28ζ CAR-T cells after the expansion from varying rounds of CAREp-stimulation by TCR repertoire analysis. (**b**) Dominant clones with more than 200 cells and their cell number distribution at different timepoints. (**c**) UMAP locations of traceable clones (clone size > 50 cells) with different proliferation capacity: group 1 (G1)−untraceable at day-30-40; group 2 (G2)−traceable at day-60-70; group 3 (G3)−traceable at day-90-100. G3-1, G3-2, and G3-3 represent group 3 cells exist at day-30-40, day-60-70, and day-90-100, respectively. (**d-f**) Balloon plot showing expression of key markers associated with memory (green line), effector (blue line), and exhaustion (orange line) phenotypes in assorted groups in **c**. **d-e** show RNA levels from GEX data, and **f** shows protein levels from CITE data. (**g**) Violin plots represent GSVA-CR and NR scores of leidens 1-3 (Fig. 5d) and G3-1. (**h**) Violin plots represent GSVA-CR, NR, and FOXO1 regulon scores of assorted groups in **c**. *P* values in **g-h** are from One-way ANOVA with Tukey’s multiple comparisons.

G3 clones were then tracked at different timepoints, and a relatively stable immune profile with a gradual differentiation shift was observed along massive expansion (**Fig. 6c**). For example, RNA levels of FAS, PTGER2, BHLHE40^46^, BATF, TOX, TIGIT, CTLA4, ENTPD1, and CD38, and protein levels of CD25, CD39, CD95 showed a gradual increase, while T cell proliferation transcription factor MYC (RNA) gradually decreased, and TNF (RNA), ITGB7 (RNA), CD69 (protein), CD134 (protein), CD45RO (protein), CD122 (protein), and CX3CR1 (protein) firstly increased then decreased (**Fig. 6d-f**). When we run gene set scoring of CR, NR, and FOXO1 regulon gene sets, G3 at 30-40 days (G3-1) showed the best score comparable to Leiden cluster-1 (**Fig. 6g**). However, cell quality indicated by CR and NR scores decreased over time (G3-2 and G3-3 compared to G3-1) (**Fig. 6h**). This suggests that by ∼day 90-100 (>8 stimulations), even the most durable clones reach a natural limit, coinciding with the plateau of expansion and cytotoxicity observed earlier (**Fig. 1, 2, 4**). In other words, while CAREp delays the onset of terminal differentiation, it does not entirely prevent it—cells eventually approach their physiological lifespan. Additionally, CAR RNA expression spectrum among clones of different longevity did not show clear difference (**Supplementary Fig. 6e,f)**. Altogether, these data demonstrate that CAREp stimulations selectively expanded durable clones and bifurcate cells to a distinct differentiation trajectory with massive proliferation and a sustained, less-exhausted phenotype, which may underlie their improved therapeutic potential.

## Conclusion and discussion

In summary, we achieved sustained in vitro expansion of human CD8⁺ CAR-T cells for over 100 days using biomaterial-mediated stimulation, revealing the intrinsic proliferative potential of human T cells under optimally structured cues^9, 11, 12^. Balanced delivery of CAR antigen and costimulatory signals enabled massive expansion not possible with tumor cells or CD3/28-Dynabeads, which led to rapid attrition. For 4-1BBζ-CARs, low-density a-CD28 costimulation ([9:1] ratio) was essential for long-term proliferation, while CD28ζ-CARs required minimal or no external costimulation in most donors. These precisely engineered external signals induced a distinct intracellular program: upregulating pathways linked to DNA repair, chromatin remodeling, telomere maintenance, and mitochondrial function, while downregulating calcium-NFAT, JAK–STAT, ERK, and cytotoxicity-associated signaling. Unlike natural stimulation that couples proliferation with exhaustion, CAREp decoupled these processes, promoting cell division without triggering terminal differentiation. These early-phase programs aligned with long-term outcomes—sustained proliferation, progenitor phenotype retention, and metabolic fitness.

Although CAREp stimulation is TCR-independent, clonal enrichment was observed across donors, indicating that the peripheral repertoire contains clones with intrinsically greater proliferative potential. CAREp consistently expanded these memory-like clones and reduced donor-to-donor variability^6, 8, 33^—suggesting a platform capable of enriching high-fitness populations often lost under standard protocols. Extensively expanded cells exhibited features of physiological regulation. Proliferation ceased at any time upon CAREp withdrawal; target cytotoxicity reduced after 11 rounds of expansion; telomerase was only transiently induced; and transcriptomic changes after ∼60–70 days indicated progression toward terminal states (increased FAS, ENTPD1, CTLA4, TIGIT, BHLHE40, BATF, TOX and decreased ITGB7, MYC, CCL5, GZMA)^46, 49^.

Interestingly, CAREp-stimulation bifurcated human T cell into durable states distinct from the normal trajectory. Morphologically, they upregulated CD25 expression and relied on external IL-2 or IL-7/IL-15 supply to proliferate. Meanwhile, the expanded cells did not secrete a high level of IFN-γ when exposed to tumor targets, although their cytotoxicity and TNF-α/IL-2 secretion were well retained or even improved after initial stimulation rounds. Considering that several studies showed that IFN-γ signaling is dispensable and even detrimental to CD8^+^ T cell expansion and memory differentiation^43–45, 51^, the reduction of IFN-γ secretion may contribute to the durable cell proliferation. Additionally, the constant high level of CD25 expression, the absence of PD-1, TIGIT, and KLRG-1 expression after extensive stimulation and expansion, as well as the coordinated high levels of TOX and STAT5, suggested unique memory-like and durable effector-like states^48^.

These durable CAR-T cell states mirror clinical predictors of persistence and efficacy^33, 36, 50^. Our GSVA analysis confirmed strong alignment with remission-associated signatures. Given the correlation between expansion quality and clinical manufacturing success, this approach may also benefit emerging allogeneic T cell therapies^52, 53^. Our previous study aiming at testing *in vivo* anti-tumor efficacy of the expanded cells faced challenges due to the inherent ‘off-tumor’ toxicity of EGFR-CAR construct, and our ongoing work focuses on affirming the therapeutic applicability of this approach for FDA-approved CAR constructs. Nevertheless, this study unprecedently demonstrates the superb expansion and functional capacities of human CAR-T cells in their physiological conditions and the signaling mechanisms imposed by the ideal external stimulation, thereby shedding light on T cell fundamentals and therapeutic engineering designs.

## Data availability

Any raw data that support the plots within this paper are available from the authors upon reasonable request.

## Acknowledgements

We acknowledge the support by the National Institutes of Health Grant 1U54CA244438, Coulter-Drexel Translational Research Partnership Program, and Drexel University Startup Fund. We thank A. Nguyen, O. Katkade, Y. Sun, T. Matteson, and A. Baetica for help on single-cell RNA-sequencing and data analysis.

## Author contributions

X.H., T.A.D., Q.T., W.A.L., C.J.Y, and J.E. conceptualized and/or supervised the study. X.H. and Q.T. designed the experiments and interpreted the results. X.H., Q.Z., L.F., L.F., Y.C., H.H., J.Z.W., R.H. performed the experiments including biomaterial engineering, T cell transduction, flow cytometry, cytotoxicity assay, and mitochondrial function analysis. L.F. and T.M. performed single-cell and bulk RNA sequencing analysis. G.A., R.A., and R.H. contributed to experimental designs and CAR construct cloning. X.H. and Q.Z. wrote the manuscript with input from all coauthors.

