## Supplemental Materials for "Programmable microparticles rewire CAR signaling to enable super-physiological expansion of human T cells *in vitro*"

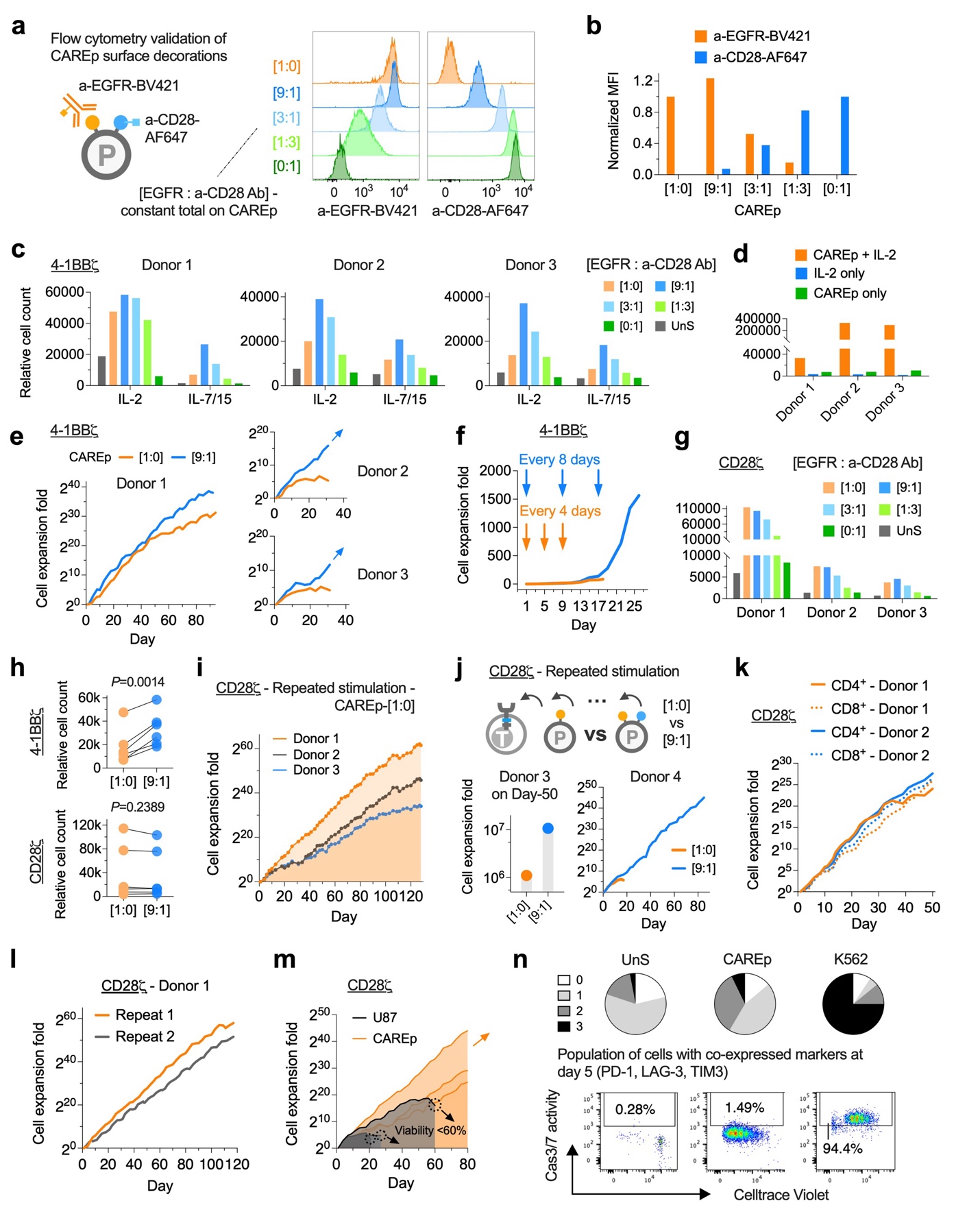

Supplementary Figure 1 CAR-T cell expansion under different stimulation conditions.

(**a**) Flow cytometry validation of CAREp surface functionalities after immunostaining. AF647-labeled a-CD28 antibodies and EGFR antigens were co-loaded on PLGA microparticles with varying densities and ratios of sequence-matched DNA-scaffolds – ([1:0], [9:1], [3:1], [1:3], [0:1]), followed by a-EGFR-BV421 staining. (**b**) Normalized mean fluorescence intensity (MFI) of fluorescently labeled a-EGFR and a-CD28 antibodies. (**c**) Relative cell count of CD8^+^ 4-1BBζ CAR-T cells at 10 days after the first stimulation by various CAREp conditions in the presence of IL-2 (30 IU/mL) or IL-7/15 (IL-7 and IL-15 at 5 ng/mL each). (**d**) Relative cell count of CD8^+^ 4-1BBζ CAR-T cells with or without IL-2 supplementation. (**e**) Cumulative expansion of CD8^+^ 4-1BBζ CAR-T cells under repeated stimulation using CAREp ([1:0] and [9:1]) every 8-10 days. (**f**) Cumulative expansion of CD8^+^ 4-1BBζ CAR-T cells under repeated CAREp ([1:0]) stimulations every 4 days or 8 days. (**g**) Relative cell count of CD8^+^ CD28ζ CAR-T cells at 10 days after the first stimulation by various CAREp conditions in the presence of IL-2 (30 IU/mL). UnS: unstimulated cells. (**h**) Relative cells count of CD8^+^ 4-1BBζ and CD28ζ CAR-T cells at 10 days after the first stimulation of CAREp conditions ([1:0] and [9:1]). Data represent n = 6 biological replicates from three human donors, and *P* values were determined by two-tailed paired *t*-test. (**i**) Cumulative expansion of CD8^+^ CD28ζ CAR-T cells under repeated stimulations by CAREp- [1:0] every 8-10 days when cells are rested from the previous stimulation. (**j**) Cumulative expansion of CD8^+^ CD28ζ CAR-T cells (engineered from donor 3 and 4) under repeated stimulations by CAREp-[1:0] versus CAREp-[9:1]. (**k**) Cumulative expansion of CD4^+^ and CD8^+^ CD28ζ CAR-T cells under repeated stimulations by CAREp-[1:0]. (**l**) Cumulative expansion of CD8^+^ CD28ζ CAR-T cells engineered from the same donor but repetitively stimulated in two batches. (**m**) Cumulative expansion of CD8^+^ CD28ζ CAR-T cells under repeated stimulations by CAREp-[1:0] versus U87 cells. Curves represent data from n = 3 donors. (**n**) Population of CD8^+^ CD28ζ CAR-T cells with co-expressed inhibitory markers (PD-1, LAG-3, TIM3) and elevated Caspase-3/7 (Cas3/7) activity after resting from one-round stimulation between CAREp- [1:0] versus EGFR-overexpressing K562 cells compared to unstimulated cells (UnS).

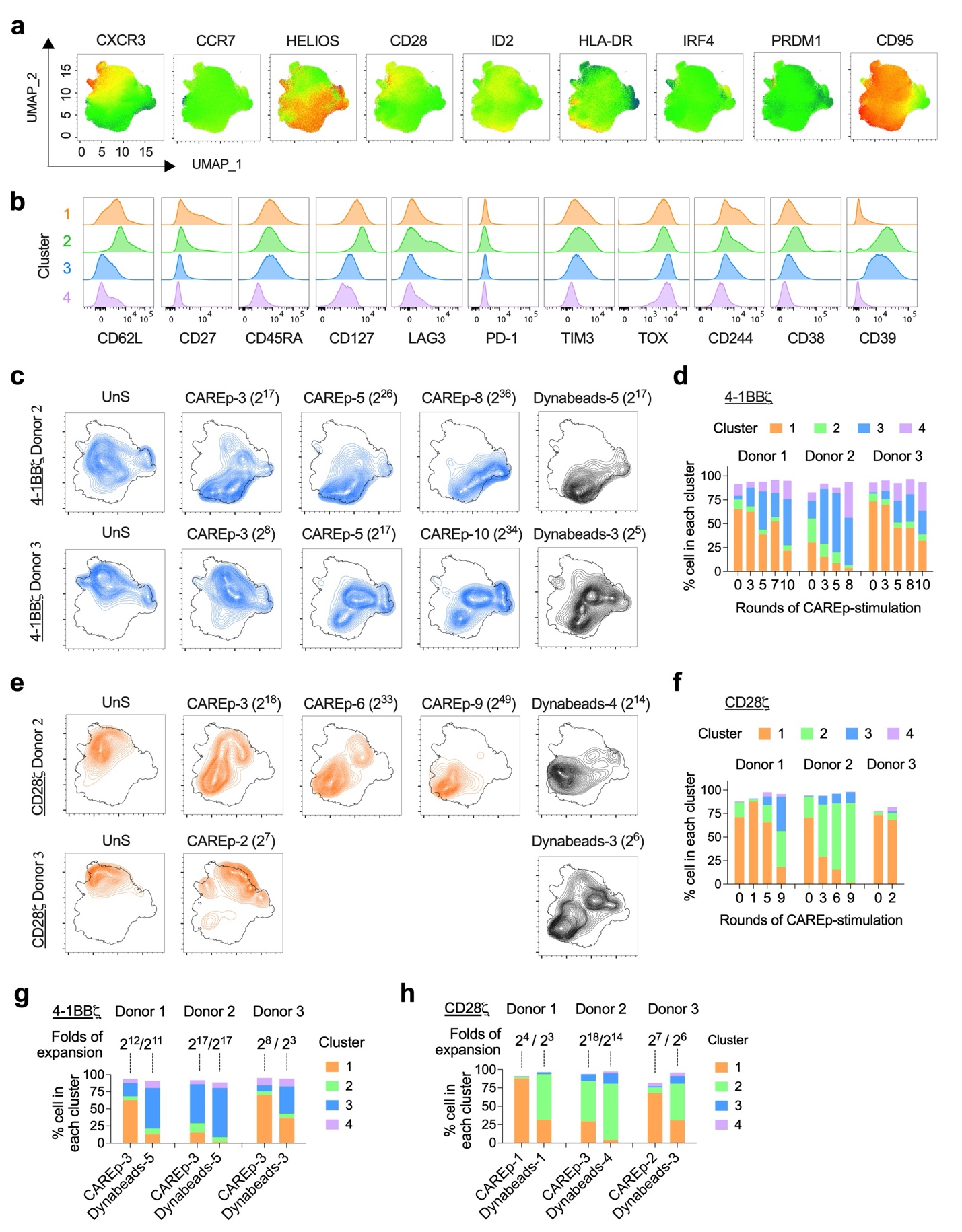

Supplementary Figure 2 CyTek immune panel profiling of CAREp or CD3/28-Dynabeads expanded CAR-T cells.

(**a**) Expression of key markers on the UMAP composed of the pooled CD8^+^ EGFR-CAR T cell samples from repeated CAREp or CD3/28-Dynabeads stimulations after the immunostaining using a 25-marker spectrum flow panel. Samples are from n = 3 donors. (**b**) Histogram of marker expression on clusters outlined in. (**c**) Distribution of the expanded 4-1BBζ CAR-T cells from CAREp or Dynabeads stimulations on the pooled UMAP in (**a**). (**d**) Population of the clusters in different rounds of CAREp stimulated 4-1BBζ CAR-T cells. (**e**) Distribution of the expanded CD28ζ CAR-T cells from CAREp or Dynabeads stimulations on the pooled UMAP in (**a**). (**f**) Population of the clusters in different rounds of CAREp stimulated CD28ζ CAR-T cells. (**g-h**) Population of the clusters in 4-1BBζ CAR-T cells (**f**) and CD28ζ CAR-T cells (**g**) with equivalent folds of expansion between varying rounds of CAREp and Dynabeads stimulations.

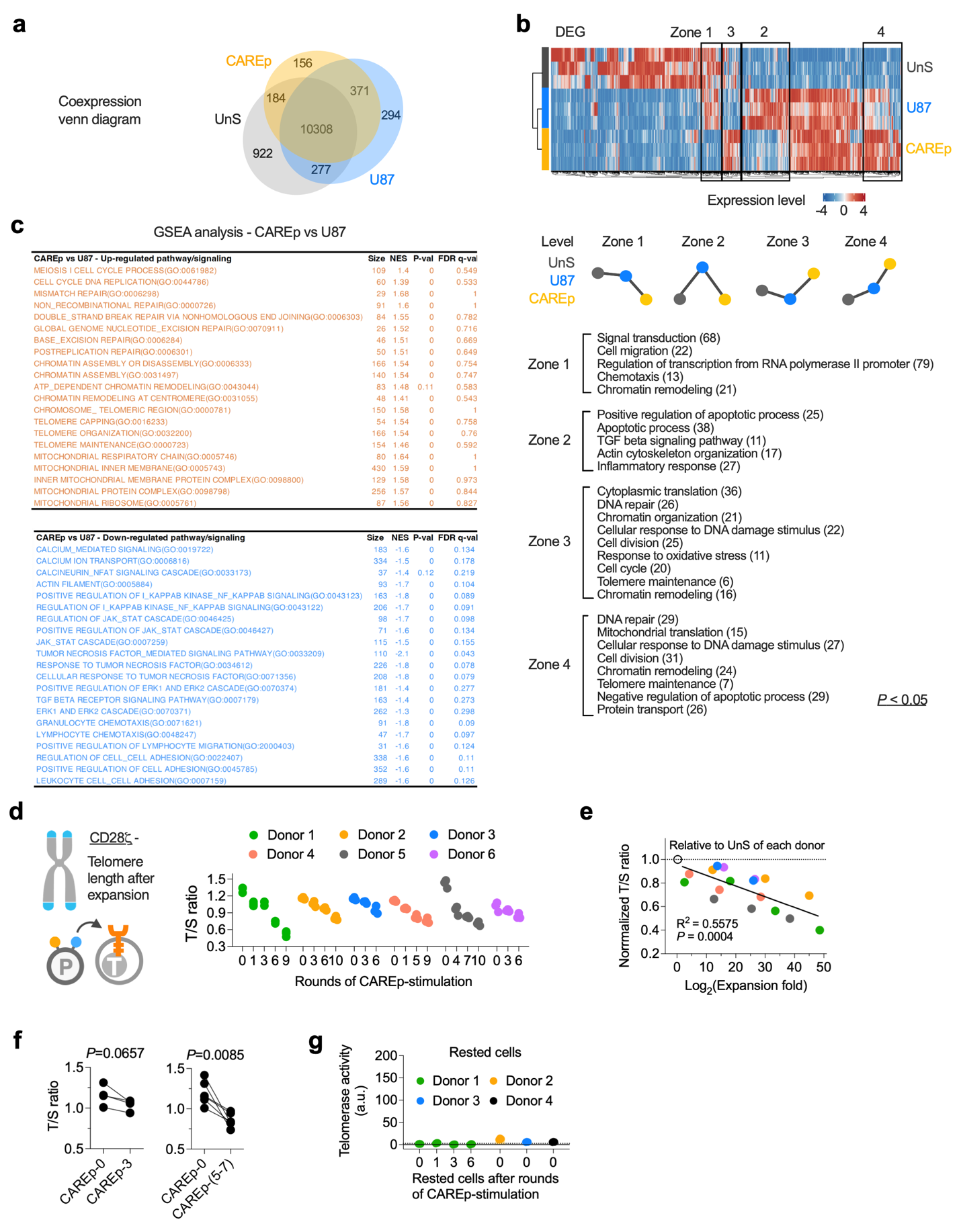

Supplementary Figure 3 Telomere length of CAREp-expanded cells and acutely induced signaling by CAREp.

(**a-b**) Coexpression venn diagram of overlapped genes (**a**) and heatmap of differentially expressed genes (**b**) from bulk RNA sequencing data of unstimulated (UnS), U87-stimulated, and CAREp-stimulated CD8^+^ CD28ζ CAR-T cells after 24 hours. Zones of interest along with associated pathways are denoted in (**b**). (**c**) Table of GSEA analysis results of various up/down regulated pathways between U87 and CAREp-stimulated cells, with normalized enrichment scores (NES) and corresponding P and Q values. (**d**) Telomere lengths of the expanded CD8^+^ CD28ζ CAR-T cells after varying rounds of CAREp stimulation by the T/S assay. Data represent mean ± SD from n = 3 technical replicates among six donors. (**e**) Linear regression between generations of cell expansion and the normalized T/S ratios to the unstimulated cells of each donor (n = 6 donors). (**f**) T/S ratio shift of cells between various rounds of stimulation. *P* values were determined by two-tailed paired *t*-test. (**g**) Telomerase activity of resting CD8^+^ CD28ζ CAR-T cells from 0, 1, 3, and 6 rounds of CAREp-expansion prior to an additional stimulation in (**f**).

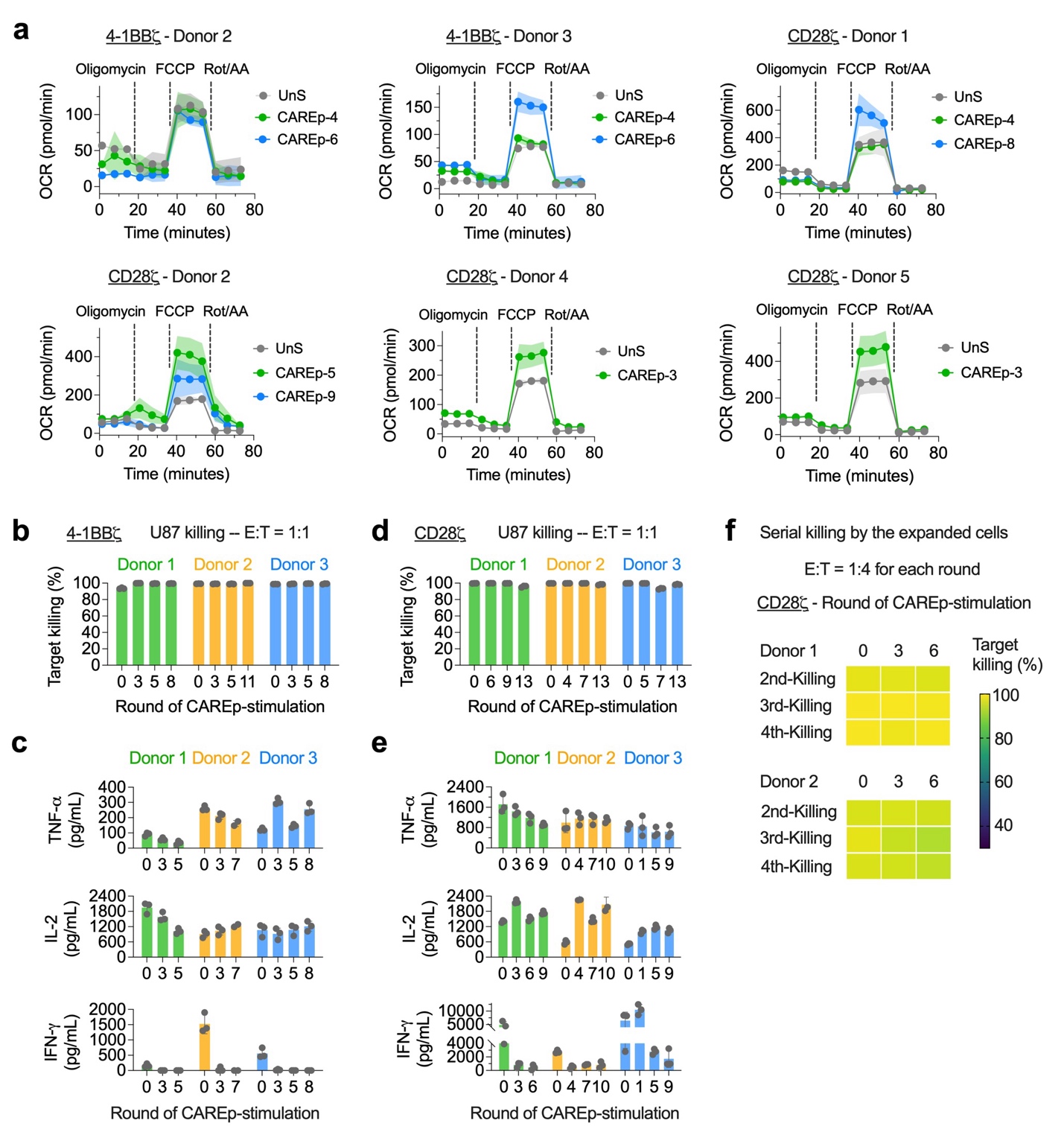

Supplementary Figure 4 Effector function of the expanded cells by Seahorse MitoStress analysis and *in vitro* target killing assay.

(**a**) Representative oxygen consumption rate (OCR) of CAREp-expanded CD8^+^ 4-1BBζ and CD28ζ CAR-T cells among donors that are treated with respiration inhibitors (Oligomycin, FCCP, Rot/AA) in the Seahorse Mitostress assay. Data represent mean ± SD from n = 6 biological replicates. (**b**) Target killing efficacy (%) of CAREp-expanded CD8^+^ 4-1BBζ CAR-T cells against U87 cell line at 72 hours after the co-culture. (**c**) TNF-α, IL-2, and IFN-γ secretion of the expanded cells at 48 hours after the co-culture with U87 cells in (**b**). (**d**) Target killing efficacy (%) of CAREp-expanded CD8^+^ CD28ζ CAR-T cells against U87 cell line at 72 hours after the co-culture. (**e**) TNF-α, IL-2, and IFN-γ secretion of the expanded cells at 48 hours after the co-culture with U87 cells in (**d**). Effector-to-target (E:T) ratio was controlled at 1:1 in (**b-e**), and data represent mean ± SD from n = 4 biological replicates. (**f**) Serial killing of CAREp-expanded CD8^+^ CD28ζ CAR-T cells against U87 cells for four rounds. E:T ratio is controlled at 1:4 for each round. Data is averaged from n = 4 biological replicates.

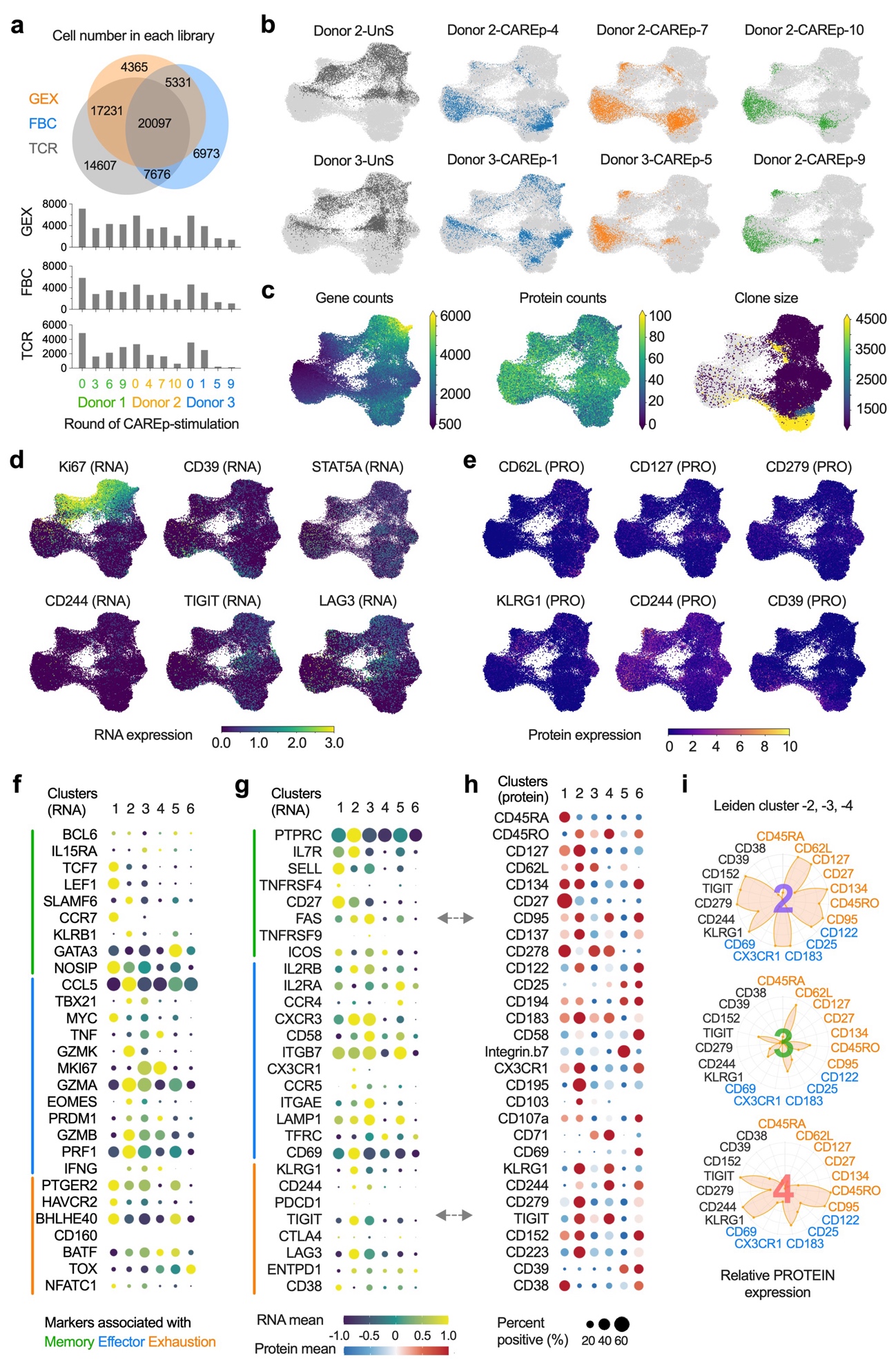

Supplementary Figure 5 Profiling of CAREp-expanded cells by single-cell RNA-sequencing.

(**a**) Size of each library from the simultaneous analysis of the transcriptome (GEX), 130 surface markers (CITE), and TCR repertoire (TCR), with the individual cell counts broken down by donor and CAREp-stimulation round. Cell-barcode overlaps between libraries are shown. (**b**) UMAPs of GEX data highlight cell distribution according to the round of CAREp-stimulation. Light grey dots represent cells from all samples. (**c**) Distribution of gene/protein/clone counts from the GEX, FBC, and TCR analyses. (**d**) Relative RNA expression of selected genes on the GEX-UMAP. (**e**) Relative protein expression of selected proteins on the GEX-UMAP. (**f-h**) Balloon plot showing expression of key markers associated with memory (green line), effector (blue line), and exhaustion (orange line) phenotypes, grouped by clusters determined via the unsupervised Leiden clustering of the GEX dataset. (**i**) Relative expression of selected surface markers (protein) in Leiden clusters 2, 3, and 4. Markers in orange, blue, and black are associated with memory, effector, and exhaustion phenotype respectively.

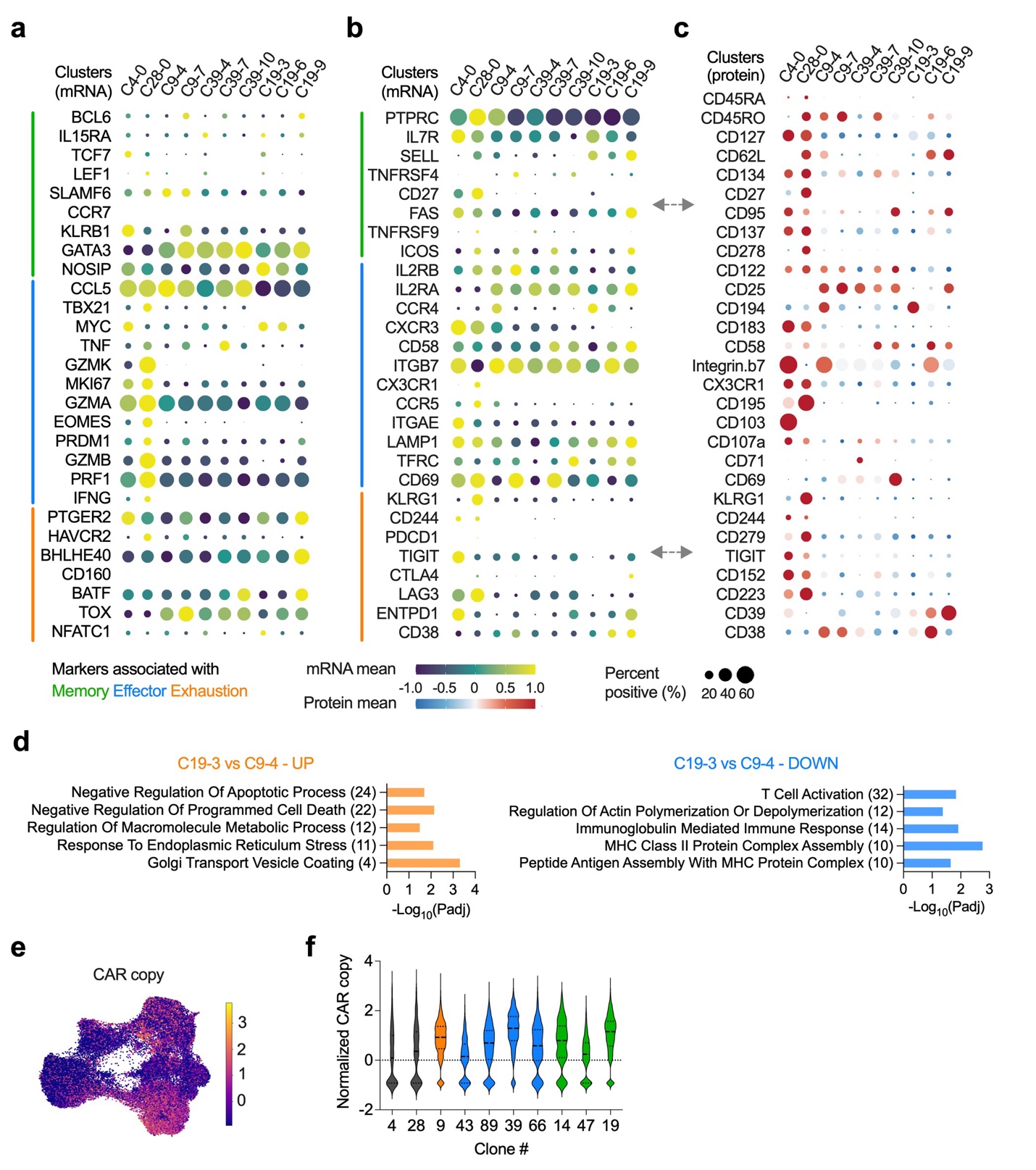

Supplementary Figure 6 Clonal tracking of the expanded CAR-T cells.

**(a-c)** Balloon plots of RNA (**a-b**) and protein (**c**) expression of key markers associated with memory (green line), effector (blue line), and exhaustion (orange line) phenotypes, grouped by clones identified from TCR data analysis along 0-9 rounds of CAREp-stimulation (**b**). (**d**) Plots displaying up/down regulated pathways between C19-3 and C9-4 clones. (**e**) CAR copy distribution displayed on UMAP. (**f**) Violin plot showing CAR copy count grouped by clone ID.

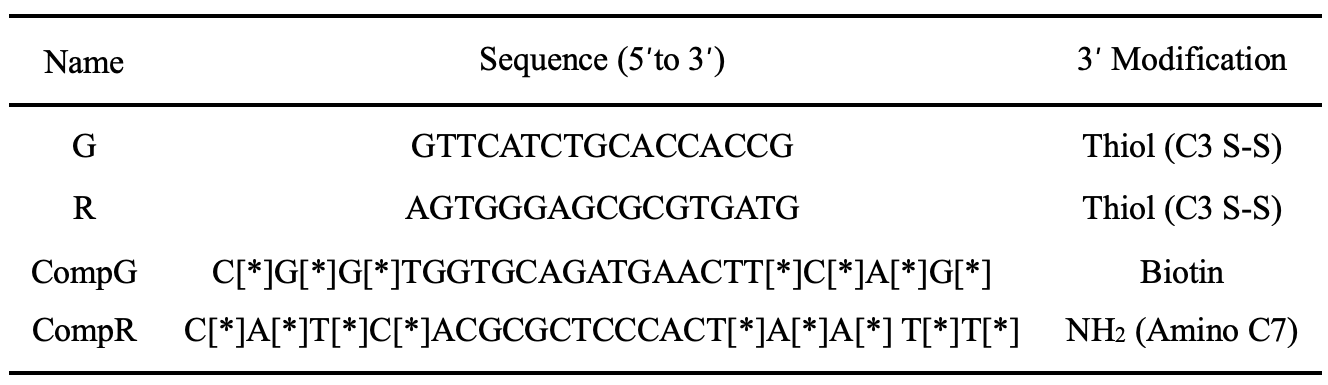

[*] Indicates an internal phosphorothioate bond.

Supplementary Table 1 DNA sequences used for CAREp surface functionalization.

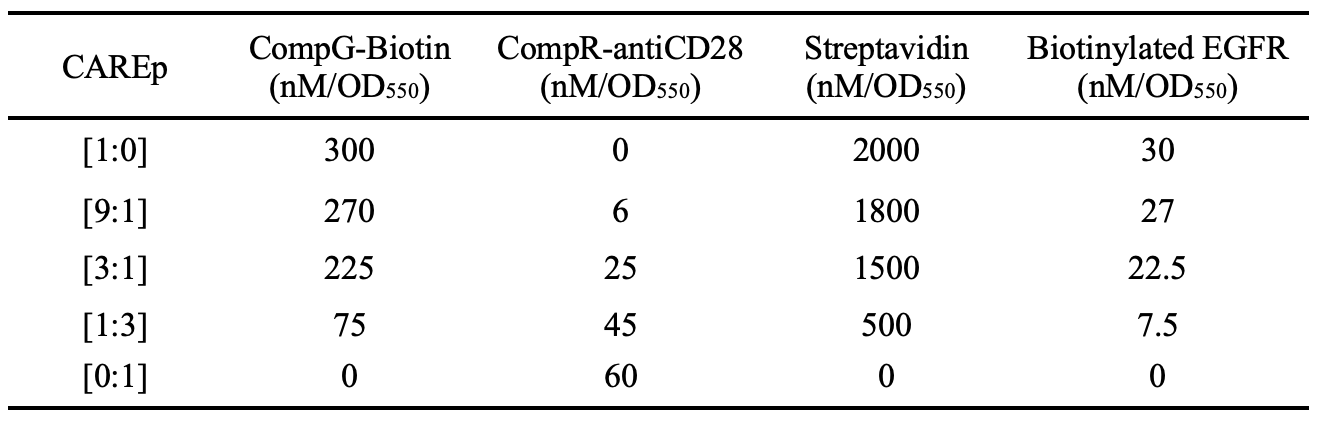

Supplementary Table 2 Molar excess of functional modules across CAREp fabrication steps.
